# Inferring amino acid interactions underlying protein function

**DOI:** 10.1101/215368

**Authors:** Victor H. Salinas, Rama Ranganathan

**Affiliations:** Green Center for Systems Biology, UT Southwestern Medical Center, 6001 Forest Park Road, Dallas, TX 75390; Departments of Biophysics and Pharmacology, UT Southwestern Medical Center, 6001 Forest Park Road, Dallas, TX 75390

## Abstract

Protein function arises from a poorly defined pattern of cooperative energetic interactions between amino acid residues. Strategies for deducing this pattern have been proposed, but lack of benchmark data has limited experimental verification. Here, we extend deep-mutation technologies to enable measurement of many thousands of pairwise amino acid couplings in members of a protein family. The data show that despite great evolutionary divergence, homologous proteins conserve a sparse, spatially distributed network of cooperative interactions between amino acids that underlies function. This pattern is quantitatively captured in the coevolution of amino acid positions, especially as indicated by the statistical coupling analysis (SCA), providing experimental confirmation of the key tenets of this method. This work establishes a clear link between physical constraints on protein function and sequence analysis, enabling a general practical approach for understanding the structural basis for protein function.

---

The basic biological properties of proteins -- structure, function, and evolvability -- arise from the pattern of energetic interactions between amino acid residues *(1–5)*. This pattern represents the foundation for defining how proteins work, for engineering new activities, and for understanding their origin through the process of evolution. However, the problem of deducing this pattern is extraordinarily difficult. Amino acids act heterogeneously and cooperatively in contributing to protein fitness, properties that are not simple, intuitive functions of the positions of atoms in atomic structures *(6)*. Indeed, the marginal stability of proteins and the subtlety of the fundamental forces make it so that many degenerate patterns of energetic interactions could be consistent with observed protein structures. The lack of knowledge of this pattern has precluded effective mechanistic models for the relationship between protein structure and function.

In principle, an experimental approach for deducing the pattern of interactions between amino acid residues is the thermodynamic double mutant cycle (7–9) (TDMC, Fig. 1A). In this method, the energetic coupling between two residues in a protein is probed by studying the effect of mutations at those positions, both singly and in combination. The idea is that if mutations *x* and *y* at positions *i* and *j,* respectively, act independently, the effect of the double mutation 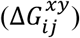 must be the sum of the effects of each single mutant 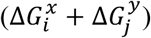. Thus, one can compute a coupling free energy between the two mutations (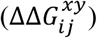 as:

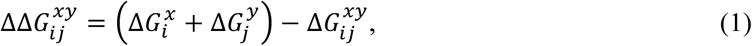

the difference between the effect predicted by the independent effects of the underlying single mutations and that of the actual double mutant. 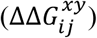 is typically proposed as an estimate for the degree of cooperativity between positions *i* and *j*.

**Figure 1:**
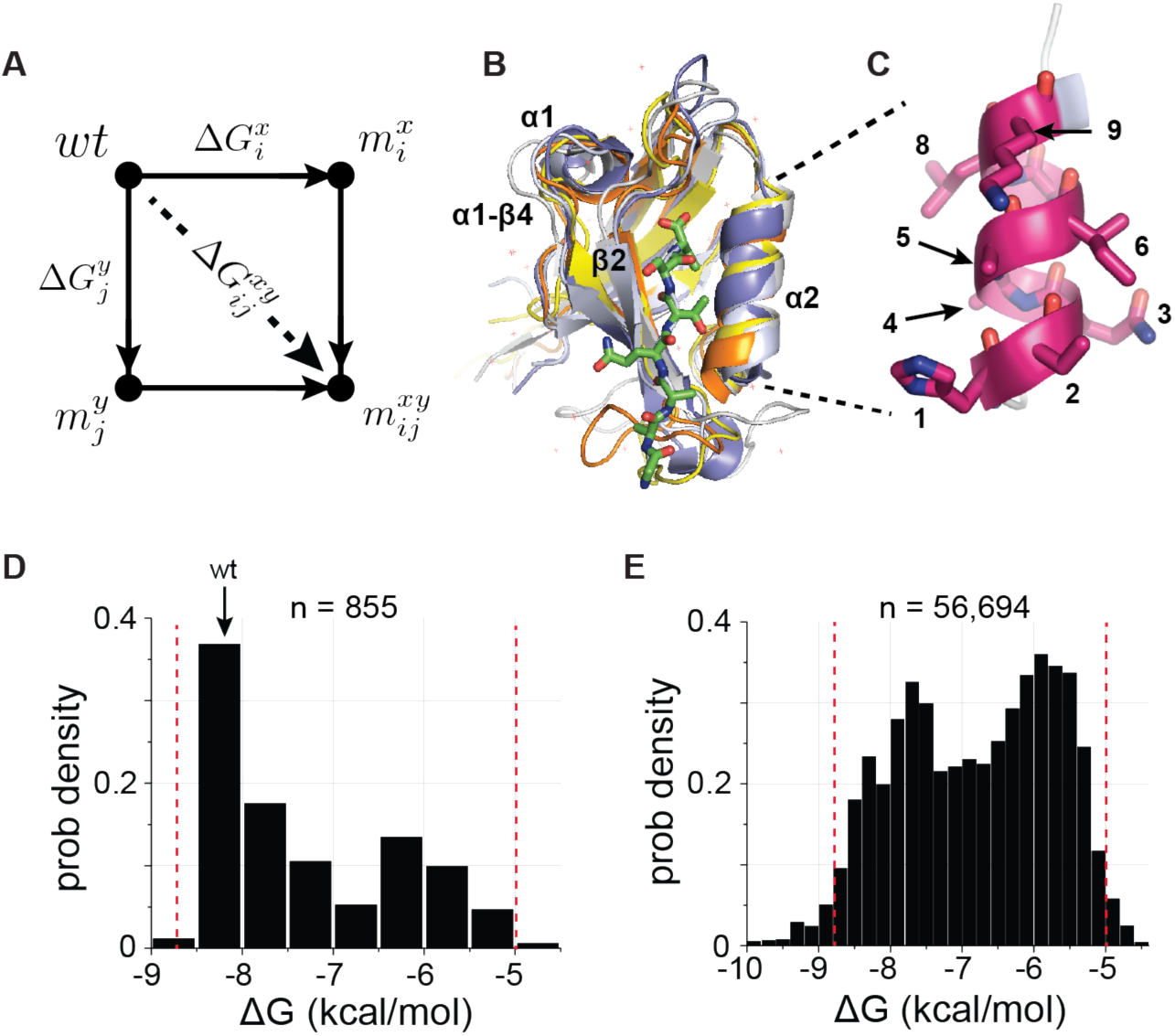
A deep coupling scan (DCS) for the PDZ binding pocket. (**A**), The thermodynamic double mutant cycle (TDMC), a formalism for studying the energetic coupling of pairs of mutations in a protein. Given two mutations (*x* at postion *i* and *y* at position *j*), the coupling free energy between them is defined as the extent to which the effect of the double mutation 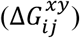 is different from the summed effect of the mutations taken individually 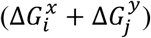, a measure of the interaction (or epistasis) between the two mutations (see Eq. 1, main text). (**B**), Structural overlay of the five PDZ homologs used in this study (PSD95^pdz3^ (1BE9, white), PSD95^pdz2^ (1QLC, orange), ZO1^pdz^ (2RRM, yellow), Shank3^pdz^ (5IZU, gray), and Syntrophin^pdz^ (1Z86, blue)), emphasizing the conserved αβ-fold architecture of these sequence-diverse proteins (33% average identity, Table S1). Structural elements discussed in this work are indicated. (**C**), The nine-amino acid α2-helix, which forms one wall of the ligand-binding site. (**D-E**), The distribution of experimentally determined binding free energies, Δ*G*_*bind*_, for all single mutations (D, 855/855) and nearly all double mutations (E, 56,694/64,980) in the α2-helix for the 5 PDZ homologs, with the affinity of wild-type PSD95^pdz3^ indicated (wt). The red lines indicate the independently validated range of the assay (Fig S1B); essentially all measurements fall within this range. These data comprise the basis for a deep analysis of conserved thermodynamic coupling in the PDZ family.

However, there are serious conceptual and technical issues with the usage of the TDMC formalism for deducing the energetic architecture of proteins. First, 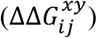 is not the coupling between the amino acids present in the wild-type protein (the “native interaction”). It is instead the energetic coupling *due to mutation,* a value that depends in complex and unknown ways on the specific choice of mutations made *(10).* Second, global application of the TDMC method requires a scale of work matched to the combinatorial complexity of all potential interactions between amino acid positions under study. For even a small protein interaction module such as the PDZ domain (~100 residues, Fig. 1B) (11), a complete pairwise analysis comprising all possible amino acid substitutions at each position involves making and quantitatively measuring the equilibrium energetic effect of nearly two million mutations. Finally, even if these two technical issues were resolved, it is unclear how to go beyond the idiosyncrasies of one particular model system to the general, system-independent constraints that underlie protein structure, function, and evolvability.

Recent technical advances in massive-scale mutagenesis of proteins open up new strategies to address all these issues. In the PDZ domain, a bacterial two-hybrid (BTH) assay for ligand-binding coupled to next-generation sequencing enables high-throughput, robust, quantitative measurement of many thousands of mutations in a single experiment - a “deep mutational scan” *(12–14).* Parameters of the BTH assay are tuned such that the binding free energy between each PDZ variant *x* and cognate ligand 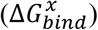 is quantitatively reported by its enrichment relative to wild-type before and after selection (Δ*E^x^*, Fig. S1 and Supplementary Methods). This relationship enables extension of single mutational scanning to very large-scale double mutant cycle analyses – a “deep coupling scan” (DCS) study *(15).* Indeed, the throughput of DCS is so high that it enables the study of double mutant cycles in several homologs of a protein family in a single experiment. Thus, DCS provides a first opportunity to deeply map the pattern and evolutionary conservation of interactions between amino acid residues in proteins, a strategy to reveal the fundamental constraints contributing to protein function.

We focused on a region of the binding pocket of the PDZ domain, a protein-interaction module that has served as a powerful model system for studying protein energetics *(13, 16).* PDZ domains are mixed αβ folds that typically recognize C-terminal peptide ligands in a binding groove formed between the α2 and β2 structural elements (Fig. 1B). We created a library of all possible single and double mutations in the nine-residue α2 helix of five sequence-diverged PDZ homologs (PSD95^pdz3^, PSD95^pdz2^, Shank3^PDZ^, Syntrophin^PDZ^, and Zo-1^PDZ^, Fig. 1C) (36 position pairs × 5 homologs, with 171 single + 12,996 double mutations + wild-type per homolog = 65,840 total variants) and measured the effect of every variant on binding its cognate ligand (Figs. 1D-E and Table S1). Independent trials of this experiment show excellent reproducibility (Fig. S2), and propagation of errors suggests an average experimental error in determining binding free energies of ~0.3 kcal/mol. Filtering for sequencing quality and counting statistics, we were able to practically collect 56,694 double mutant cycles (87% of total) for the α2 helix for all five homologs, with an average of 315 cycles per position pair per homolog (Table S1). Thus, we can (1) analyze the distributions of double mutant cycle coupling energies for nearly all pairs of mutations in the α2 helix and (2) study the divergence and conservation of these couplings over the five homologs.

We first addressed the problem of how to estimate native coupling energies from mutant cycle data. In general, the effect of a mutation at any site in a protein is a complex perturbation of the elementary forces acting between atoms, with a net effect that depends on the residue eliminated, the residue introduced, and on any associated propagated structural effects. Thus, the distribution of thermodynamic couplings at any pair of positions over many mutation pairs could in principle be arbitrary and difficult to interpret. However, we find surprising simplicity in the histograms of coupling energies. In general, the data follow a double-Gaussian distribution, with either both mean values centered at zero or with one of the means different from zero (Fig. 2, and Figs. S3-S8). Indeed, most distributions are centered close to zero, with just a few position pairs displaying two distinct populations. This result holds not only for the individual PDZ homologs selected (Figs. S3-S7) and for the average over homologs (Fig. 2), but also for DCS in an unrelated protein (GB1, Fig. S8, and *(15)),* suggesting that the binary character of couplings may be universal in proteins. A simple mechanistic model is that the observed free energy of ligand binding arises from a cooperative internal equilibrium between two functionally distinct conformational states, with mutations at some sites capable of dramatically perturbing this equilibrium (Fig. S9). Indeed, such a two-state internal equilibrium has been observed in PDZ domains, and is part of an allosteric regulatory mechanism controlling ligand binding *(17).* Thus, the population-weighted mean of the distribution of coupling energies for each position pair (Fig. 2, dashed lines) provides an empirical estimate of the native interaction between amino acids through mutagenesis.

**Figure 2:**
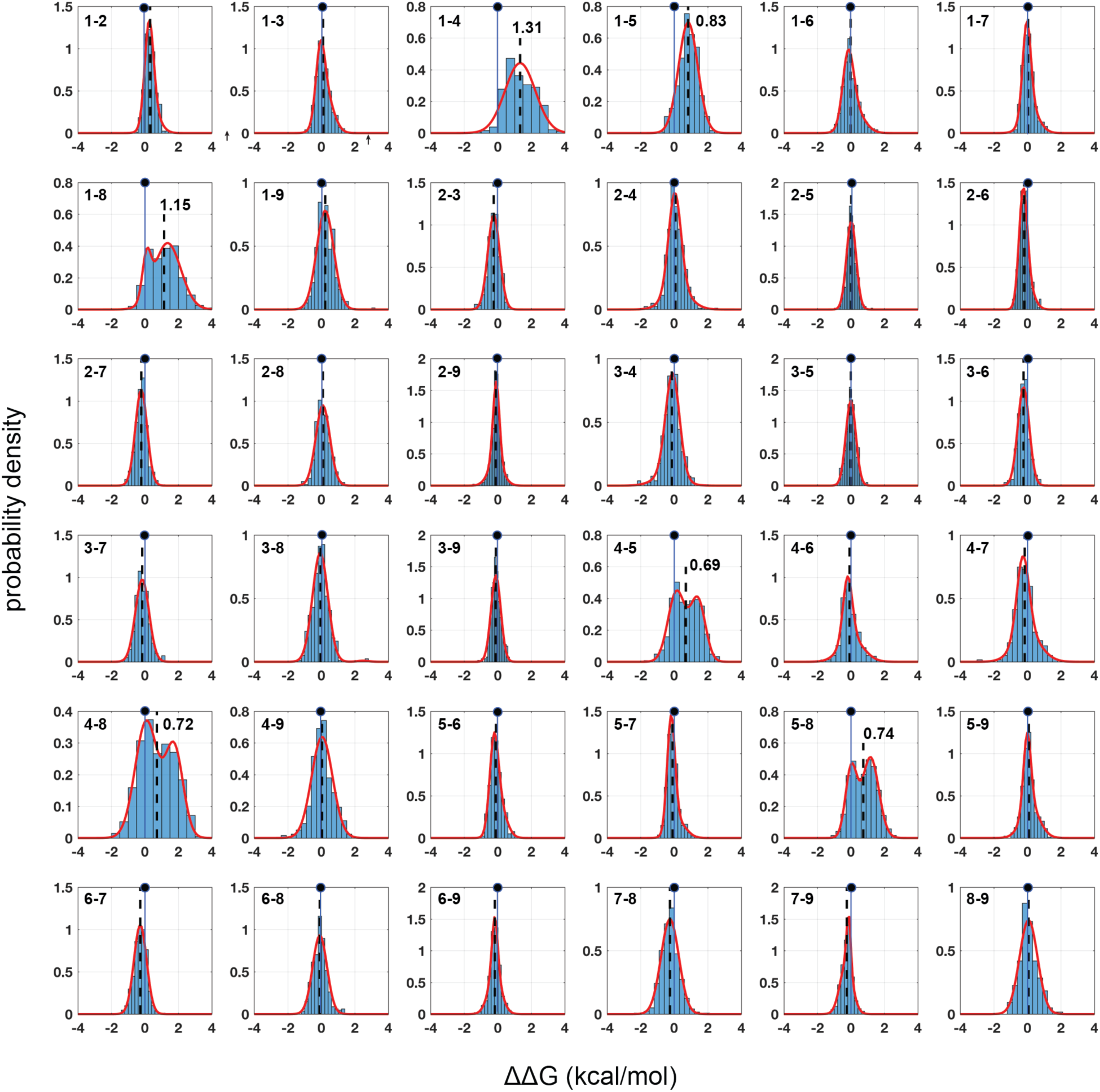
Homolog-averaged pairwise thermodynamic couplings in the PDZ domain. Each subplot shows the distribution of coupling free energies (ΔΔG, see Eq. 1, main text) for all measured mutants at one pair of positions in the α2-helix (numbering per Fig. 1C) averaged over the five homologs. The distributions are fit to single or double Gaussians, using the Bayes Information Criterion to justify choice of model, and the position of zero coupling is indicated by the solid line and circle above. Population-weighted mean values are represented by dashed lines. The data are remarkably well defined by the fitted models. Most position pairs have distributions centered close to zero, with only six pairs comprising all pairwise couplings between positions 1, 4, 5, and 8 showing deviations. For these pairs, distributions of mutational coupling follow either a single mode (1-4, 1-5) or two modes with one centered at zero (1-8, 4-5, 4-8, 5-8); population-weighted mean values for these pairs are indicated. Figures S3-S7 show these data for each homolog taken individually.

Two technical points are worth noting. First, the spread of the distributions is large, generally exceeding the estimated magnitude of the native interactions (Fig. 2). This means (1) that traditional mutant cycle studies carried out with specific choices of mutations are more likely to just reflect the choice of mutations rather than the native interaction, and (2) that the only way to obtain good estimates of the native interaction between residues is to average over the effect of many double mutant cycles per position pair. Second, we find that the BTH/sequencing approach displays such good reproducibility that it is possible to detect coupling energies with an accuracy that is on par with the best biochemical assays. For example, the average standard deviation in mean coupling energies for position pairs over four independent experimental replicates in PSD95^pdz3^ is ~0.06 kcal/mol. Thus, we can map native amino acid interactions with high-throughput without sacrificing quality.

What do the data tell us about the pattern of amino acid interactions? Figure 3 shows heat maps of the estimated native coupling energies between all pairs of amino acids within the α2 helix for each PDZ homolog. The data demonstrate both idiosyncrasy and conservation of amino acid couplings in paralogs of a protein family. For example, helix positions 3-4 show moderate couplings in two of the domains (PSD95^pdz3^ and Syntrophin^PDZ^, Fig. 3a and 3d) but not in the other homologs. Similarly, coupling between positions 7-8 is shared by PSD95^pdz3^, PSD95^pdz2^, and Zo1^PDZ^ (Figs. 3A, 3B, and 3E) but not in the other two homologs. In contrast, all pairwise interactions between positions 1, 4, 5, and 8 show a systematic pattern of energetic coupling in all homologs tested. Thus, each PDZ domain displays variations in the pattern and strength of amino acid energetic couplings, but also includes a set of evolutionarily conserved couplings at a few positions. We take the conserved couplings to represent the most fundamental constraints underlying PDZ function, with the homolog-specific couplings indicating more specialized or even serendipitous couplings.

**Figure 3:**
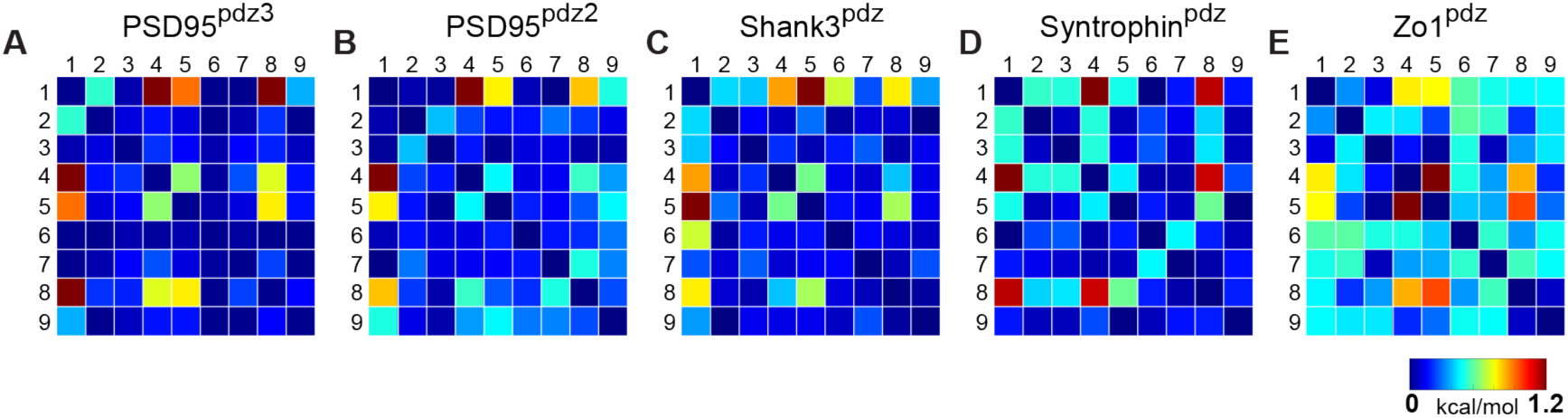
Conservation and idiosyncrasy in the pattern of energetic couplings over PDZ homologs. Matrices of mutation averaged pairwise thermodynamic couplings for the α2-helix in each PDZ homolog. The color scale is chosen to represent the full range of measured energetic couplings. The data show that some couplings are specific to individual homologs or shared by a subset of homologs, but that couplings between positions 1, 4, 5, and 8 are conserved over homologs.

To isolate the fundamental couplings, we averaged all the double mutant cycle data over all mutations and over the five PDZ homologs tested, resulting in a matrix of evolutionarily conserved pairwise thermodynamic couplings (Fig. 4A). This analysis reinforces the result that positions 1, 4, 5, and 8 comprise a cooperative network of cooperative functional positions in the PDZ domain family, and the remainder, even if in direct contact with each other or with ligand, contribute less and interact idiosyncratically. The conserved couplings form a chain of physically contiguous residues in the tertiary structure that both contact (1, 5, 8) and do not contact (4) the ligand (Fig. 4B). Interestingly, position 4 is part of a distributed allosteric mechanism in some PDZ domains for regulating ligand binding *(17),* providing a biological role for its energetic connectivity with binding pocket residues. Overall, the pattern of couplings does not just recapitulate all tertiary contacts between residues (compare Fig. 4A with 4E, white and black circles) or the pattern of internal backbone hydrogen bonds that define this secondary structure element. Instead, conserved amino acid interactions in the PDZ α2 helix are organized into a spatially inhomogeneous, cooperative network that underlies ligand binding and allosteric coupling.

**Figure 4:**
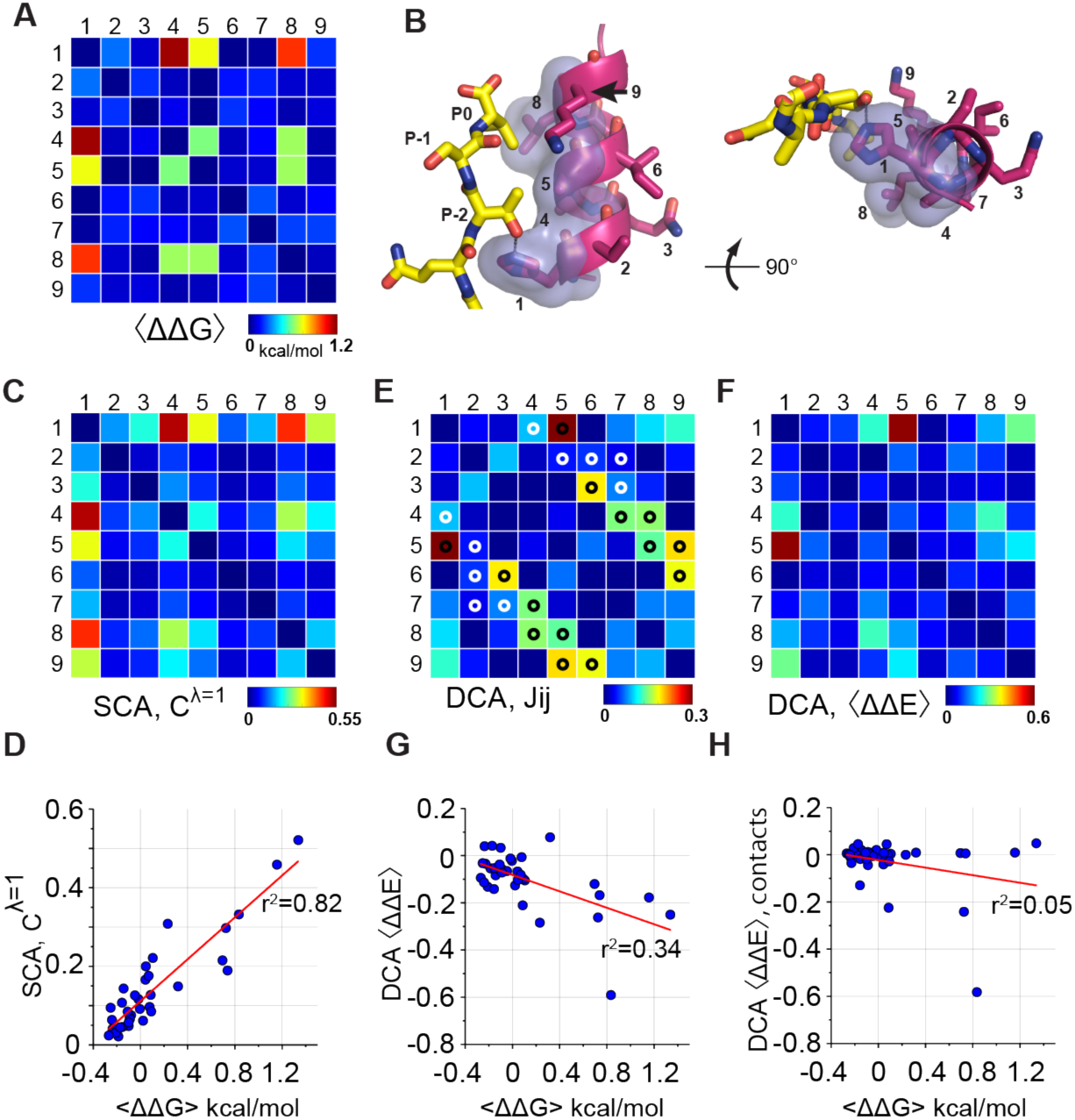
Coevolution-based inference of energetic couplings. (**A**), The homolog and mutation averaged couplings (corresponding to Fig. 2), displaying the evolutionarily conserved relationships between amino acids in the PDZ α2-helix. (**B**), Two views of the α2-helix, with positions engaged in conserved coupling in transparent surface representation, and ligand in yellow stick bonds. These include three positions in direct contact with ligand (1, 5, 8) and one allosteric position buried in the core of the protein (4). (**C**), Coevolution of sequence positions corresponding to the top eigenmode of the SCA matrix, derived from an alignment of 1689 eukaryotic PDZ domains. The data show that a subset of positions coevolve within the PDZ α2-helix. (**D**), The relationship between experimental homology-averaged energetic couplings (<ΔΔG>) and SCA-based coevolution. (**E**), The matrix of direct couplings (*Jij*) from the DCA method, with tertiary contacts in the PDZ structure (1BE9) indicated by white or black circles; by convention *(25),* trivial contacts between residues with sequence distance less than three are not shown. The data show that all top direct couplings identified by DCA are indeed tertiary structural contacts. (**F**), Mutation-and homolog-averaged energetic couplings inferred from the DCA model, computed precisely as for the experimental data in panel B; see Supplementary Methods for details. (**G**), The relationship between experimental (<ΔΔG>) and DCA-inferred (<ΔΔE>) couplings in the PDZ α2-helix. (**H**), The relationship between experimental and DCA-inferred couplings from in which top couplings defining contacts are preserved and all non-contact couplings are randomly scrambled. The data show that pairwise couplings in the DCA model between non-contacting positions contribute significantly to prediction of protein function.

This result begins to expose the complex energetic couplings underlying protein function, but also highlights the massive scale of experiments required to deduce this information for even a few amino acid positions. How can we generalize this analysis to deduce all amino acid interactions in a protein, and for many different proteins? There are potential strategies for pushing deep mutational coupling to larger scale, but quantitative assays such as the BTH are difficult to develop, mutation libraries grow exponentially with protein size, and the averaging over homologs will always be laborious, expensive, and incomplete.

A different approach is suggested by understanding the rules learned in this experimental study for discovering relevant energetic interactions within proteins. The bottom line is the need to apply two kinds of averaging. Averaging over many mutations provides an estimate of native interaction energies between positions, and averaging the mutational effects over an ensemble of homologs separates the idiosyncrasies of individual proteins from that which is conserved in the protein family. Interestingly, these same rules also comprise the philosophical basis for a class of methods for estimating amino acid couplings through statistical analysis of protein sequences. The central premise is that the relevant energetic coupling of two residues in a protein should be reflected in the correlated evolution (coevolution) of those positions in sequences comprising a protein family *(16, 18–20).* Statistical coevolution also represents a kind of combined averaging over mutations and homologs, and if experimentally verified, would (unlike deep mutational studies) represent a scalable and general approach for learning the architecture of amino acid interactions underlying function in a protein. The data collected here provides the first benchmark data to deeply test the predictive power of coevolution-based methods.

One approach for coevolution is the statistical coupling analysis (SCA), a method based on measuring the conservation-weighted correlation of positions in a multiple sequence alignment, with the idea that these represent the relevant couplings *(16, 21).* In the PDZ domain family (~1600 sequences, pySCA6.0 (22)), SCA reveals a sparse internal organization in which most positions evolve in a nearly independent manner and a few (~20%) are engaged in a pattern of mutual coevolution *(16, 21, 22)*. In this case, the coevolving positions are simply defined by the top eigenmode (or principal component) of the SCA coevolution matrix, and represent a biologically important allosteric mechanism connecting the β2-β3 loop with the α1-β4 surface through the binding pocket and the buried α1 helix (Fig. 1B, and *(23)).* Extracting the coevolution pattern in the top eigenmode for just the α2 helix (Fig. 4C), we find that coevolution as defined by SCA in fact nearly quantitatively recapitulates the homolog-averaged experimental couplings collected here (*r*^2^ = 0.82, *p* = 10^−14^ by F-test, Fig. 4D). The predictions also hold for individual homologs (Fig. S10A-E), consistent with the premise that the essential physical constraints underlying function are deeply conserved. This relationship is robust to alignment size and method of construction (Fig. S11A-C), but depends on both of the basic tenets that underlie the SCA method – conservation-weighting (Fig. S11D-E) and correlation (Fig. S11 F-G) (22).

Another approach for amino acid coevolution is direct contact analysis (DCA, *(24, 25)),* a method developed for the prediction of tertiary contacts in protein structures. DCA uses classical methods in statistical physics to deduce a matrix of minimal pairwise couplings between positions (*J*_*ij*_, Fig. 4E) that can account for the observed correlations between amino acids in a protein alignment, with the hypothesis that the strong couplings in *j*_*ij*_ will be direct contacts in the tertiary structure. Indeed, studies convincingly demonstrate that the top *L/2* (where L is the length of the protein) couplings are highly enriched in direct structural contacts (26). Consistent with this, this method successfully identifies direct contacts in the PDZ α2 helix (Fig. 4E, compare heat map to white and black circles) to an extent that agrees with the reported work. However, DCA-based predictions of functional energetic couplings between mutations are weakly (though significantly) related to the homolog-averaged experimental data (*r*^2^ = 0.33, *p* = 10^−3^ by F-test, Fig. 4F-G). These results are similar or poorer for prediction of couplings in individual domains (Fig. S10G-K). Interestingly, a DCA model in which only the top pairwise couplings in *J*_*ij*_ that define tertiary contacts are retained and the weaker non-contacting couplings are randomly scrambled shows predictions that are unrelated to the experimental data (*r*^2^ = 0.05, *p* = 0.09 by F-test, Fig. 4H). Thus, non-contact couplings in the DCA model, which represent noise from the point of view of structure prediction, contribute significantly to prediction of function.

These findings clarify the current state of sequence-based inference of protein structure and function *(27, 28).* DCA successfully predicts contacts in protein structures in the top couplings, but in its current form, does not appear to capture the cooperative constraints that underlie protein function well. In contrast, SCA does not predict direct structural contacts well, but instead seems to more accurately capture the energetic couplings that contribute to protein function(s). As explained previously, these two approaches sample different parts of the information contained in a sequence alignment *(29, 30),* and therefore are not mutually incompatible. These results highlight the need to unify the mathematical principles of contact prediction and SCA-based energetic predictions towards a more complete model of information content in protein sequences.

In summary, the collection of functional data for some 56,000 mutations in a sampling of PDZ homologs demonstrates an evolutionarily conserved pattern of amino acid cooperativity underlying function. This pattern is well-estimated by statistical coevolution based methods, suggesting a powerful and (given the scale of experiments necessary) uniquely practical approach for mapping the architecture of couplings between amino acids. Indeed, the remarkable conclusion is that with a sufficient ensemble of sequences comprising the evolutionary history of a protein family, the pattern of relevant amino acid interactions can be inferred without any experiments.

## Acknowledgements

We thank F.J. Poelwijk, M.A. Stiffler, C. Nizak, O. Rivoire, M. Weigt, R. Monasson, and S. Cocco for discussions and technical advice, and members of the Ranganathan laboratory for critical review of the manuscript. We also thank the High Performance Computing Group (BioHPC) and the Genomics Core at UT Southwestern for providing computational resources and sequencing, respectively. This work was supported by NIH Grant RO1GM12345 (to R.R.), a Robert A. Welch Foundation Grant I-1366 (to R.R.), and the Green Center for Systems Biology at UT Southwestern Medical Center. V.S. was supported in part through a pre-doctoral fellowship (NIGMS T32 GM008203).

## Supplementary Materials

Materials and Methods

Figs. S1-S11

Table S1

Additional Data

